# Parasite-mediated predation determines the infection in a complex predator-prey system

**DOI:** 10.1101/2023.10.29.564596

**Authors:** Ana C. Híjar-Islas, Amy Milne, Christophe Eizaguirre, Weini Huang

**Author notes:** These authors contributed equally to this work.

## Abstract

The interplay of host-parasite and predator-prey interactions is critical in ecological dynamics because both predators and parasites play an important role in regulating populations and communities. But what is the prevalence of infected prey and predators when a parasite is transmitted through trophic interactions, particularly when stochastic fluctuations of demographical changes are allowed arising from individual-level dynamics? Here, we analysed the system stability and the frequency of infected and uninfected host subpopulations in a complex predator-prey-parasite system, where infection happens through trophic interactions transmitting parasites from prey to predators. We varied the parasite virulence implemented as reproductive costs imposed on infected hosts and the probabilities of parasites infecting the hosts per encounter, to investigate how those important evolutionary factors will determine the species coexistence and population composition. We further explored the role of stochasticity in our system by comparing our deterministic analysis with stochastic simulations. Our results show that parasites go extinct when the infection probabilities of either host are small. The success in infecting the final host (the predator) is more critical for the survival of the parasite species, as the threshold for infection probability of the predator is higher than that of the prey for three-species coexistence. While our stochastic simulations agree with deterministic predictions well in most parameter regions. However, in the border parameter regions between coexistence and extinction typically with high infection probabilities, while only one possible outcome in deterministic dynamics, both coexistence and extinction can happen in stochastic repeats under the same parameter values. This illustrates the importance of stochasticity and demographic fluctuations in species coexistence. In addition, the proportion of infected individuals increases with the infection probabilities in our deterministic analysis and stochastic simulations as expected. Interestingly, we found that in some parameter space, the relative frequencies of infected and uninfected individuals are different between the intermediate host (prey) and the final host (predator) populations. This counterintuitive observation shows that the interplay of host-parasite and predator-prey interactions lead to more complex dynamics than a simple resource-consumer relationship.

## 1 Introduction

Food webs are complex networks composed of species and governed by their interactions, whereby consumers feed on resources to gain energy [1, 2]. When a species goes extinct, its local network may also disappear from the food web, potentially altering the system’s stability and dynamics [3]. While food webs tend to be robust to random losses, some species may play crucial roles in maintaining the food web integrity and therefore are likely to cause the greatest damage if removed [4]. The degree of impact caused by the loss of a species is characterized by its number of interactions with other species and the degree of specialization of those interactions [3].

Parasites represent large biomass on earth [5] and are involved in an estimated 75% of food web links [6]. Parasites impact their host population through changes in body condition [7], behaviour [8], and reproduction [9]. The consequences of infection go beyond a single population because hosts are connected with other species, such as predators and prey [10]. In natural systems, parasites can infect multiple hosts directly or via trophic interactions [11]. Trophically transmitted parasites provide natural biological indicators of trophic links because they are the accumulated consequence of longterm feeding by their hosts [12, 13]. Some trophically transmitted parasites can even manipulate the behaviour of their intermediate prey host in ways that increase consumption rates, and therefore transmission to their final predator host [14, 15]. This flux transfer has consequences on the food web connectance and stability [16]. Despite their ecological importance, however, the mechanisms driving the dynamics of trophically transmitted parasites are not well understood [13].

From an ecological perspective, biological populations exhibit a large spectrum of dynamical behaviour; from stable equilibrium points, to stable cyclic oscillations, to chaotic dynamics [17].The latter can refer to the appearance of aperiodic cycles in population dynamics, and may occur when the per capita rate of increase exceeds some threshold value [18]. A biological system is considered internally stable if it does not experience significant changes in its characteristics and returns to a steady state after a perturbation [17]. Theoretical work on a complex food web model demonstrates that the fluctuations in predator-prey population dynamics change according to the attack rate of the predator [19]. Their model shows that too inefficient or too aggressive predators result in vigorous population fluctuations and prevent predator-prey coexistence [19]. Further theoretical work shows that the inclusion of parasites in predator-prey systems often leads to chaotic dynamics [20, 21, 22]. This is because epidemiological processes (such as disease) can alter the death and birth rates of the predator and prey hosts and lead to aperiodic cycles in predator-prey dynamics.

Beside classical predator-prey cycles often referred to as total abundances in each species, subpopulations of infected and uninfected hosts coexist at equilibrium [23]. The relative frequencies of infected and uninfected hosts are likely to be relevant in community dynamics. For instance, a high frequency of infected prey may have a bottom-up effect on the food web because the reduction of prey species is followed by the population decline of its consumers [24]. Also, a high frequency of infected prey increases the probability of parasite transmission to higher trophic levels. As a consequence, a high frequency of infected predators may lead to top-down effects because the reduction of predator abundance relaxes the predation pressures on the prey, allowing the prey to reproduce more rapidly [24]. The resulting increase in prey abundance leads to stronger pressures on lower trophic levels such as smaller prey or plants. These cascading effects can impact community structure and dynamics [25] and ecosystem functioning [25, 26].

Here, we built an individual-based model to capture such a complex predator-prey-parasite system, based on microscopic events, including reproduction, intrinsic death, competition, infection, and predation for relevant species. Note that when applying individual-based modelling in ecological systems, we refer to individuals as microscopic components and populations and communities as macroscopic organisational levels, hence microscopic events refer to individual-level events [27]. In our model, the parasite is transmitted trophically from infected prey to predators. We investigated the extent to which different factors such as the infection probabilities and parasite virulence (hereby defined by the reproductive costs imposed on infected hosts), affect the transmission and persistence of the trophically transmitted parasite. We decomposed the predator and prey populations into infected and uninfected subpopulations to identify the parameters that influence the stable states among the three species.

While using a stochastic individual-based model, we also wrote down the rate equations of microscopic reactions to analyse the average dynamics of our system, which resembles the population-level models like classical generalised Lotka-Volterra equations [28, 29]. Although stochastic dynamics normally agrees with its corresponding deterministic system in large populations, it allows random demographic fluctuations which are important in some evolutionary scenarios, especially for the extinctions of small populations [30]. Since antagonistic interactions such as those between parasites and hosts, and prey and predators, often lead to cyclic dynamics where population sizes could drop and go through deep valleys, we are interested in how stochasticity arising from the individual level will impact the coexistence of the three species and their population composition (i.e., frequencies of infected and uninfected host individuals).

## 2 Methods

We consider a system with three species, prey, predator and parasite denoted by *X, Y*, and *Z* respectively in our microscopic reactions, which represent individual level events like reproduction, death, competition, infection and predation. Since the parasite will infect an intermediate host, the prey, before being transmitted trophically to the definitive host, the predator, there are two distinct subtypes within each host species, the infected and uninfected. We denote the infected and uninfected prey populations by *X*_*I*_ and *X*_*U*_, respectively, and the infected and uninfected predator populations by *Y*_*I*_ and *Y*_*U*_, respectively. While reactions and their rates are defined in the individual level, how often e.g. reproduction or predation happen in the population level will also depend on the population abundances. For example, predation takes place in the predation rate per individual encounter multiplied by the prey and predator abundance. In the absence of the parasite, the prey and predator populations interactions are described by a damped Lotka-Volterra model [31, 32, 33].

### 2.1 Microscopic reactions of individuals from the three species

In the absence of the other species the prey populations can be described by the following reactions, where *g*_*x*_ ∈ ℝ ^+^ is the reproduction rate of the prey, *r*_*x*_∈ [0, 1] is the suppression of an infected preys reproduction and *K* ∈ ℝ ^+^ is the environmental carrying capacity of the prey population. While we already apply a cost of infection in reproduction, we assume that infection does not influence the competition outcome between infected and uninfected prey. However, this cost of competition can be easily implemented if necessary. We note here that infected prey offspring are uninfected.

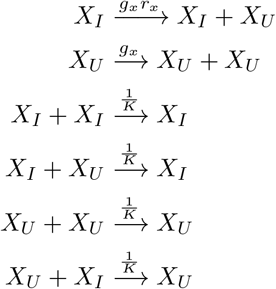

An uninfected prey encounters a free-living parasite at rate *S*, where *S* ∈ ℝ ^+^. It is then infected with probability *Q*_*x*_ ∈ [0, 1]. When a parasite successfully infects the prey, the parasite is removed from the free-living parasite population. Otherwise, it remains in free-living parasite population.

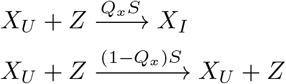

The predation rate per individual encounter between a prey and a predator is *f*_*y*_. The reproduction efficiency of the predator by consuming is *k*_*y*_ ∈ [0, 1]. When an uninfected prey meets an uninfected predator, we have the following reactions.

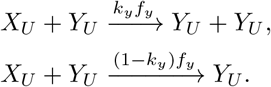

However when an infected prey meets an uninfected predator, the predator can be infected with probability *Q*_*y*_ ∈ [0, 1]. If the predator is successfully infected by the parasite through consumption of infected prey, the infected predator’s ability to reproduce is affected by the presence of the parasite and hence is suppressed according to parameter *r*_*p*_ ∈ [0, 1]. Meanwhile, the parasite transmitted from the primary host, the prey, will reproduce within the definitive host, the predator, and have *n*_*z*_ ∈ ℤ ^+^ offspring per successful infection.

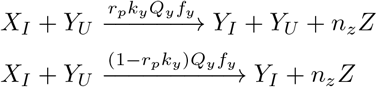

If the parasite is not successful in infecting the predator through trophic transmission, there is still a suppress of predator reproduction due to exposure *r*_*e*_ ∈ [0, 1]. We assume that the reproduction suppression due to infection is larger than the one due to exposure. Hence, *r*_*p*_ ≤ *r*_*e*_.

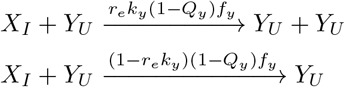

Similarly, when an uninfected prey meets an infected predator, the infected predator can reproduce with the cost *r*_*p*_. Note parasite reproduction already happened when this predator was first time infected, thus is not redundantly included here.

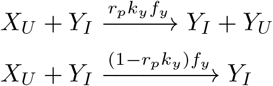

Finally, we consider the remaining possible reactions where an infected prey meets an infected predator. If the infected predator is successfully infected again by the new encountered infected prey, its ability to reproduce is further suppressed by the same parameter *r*_*p*_. The parasite transmitted by this new encountered prey will reproduce in its final host, because this is the first time this predator infected by this individual parasite.

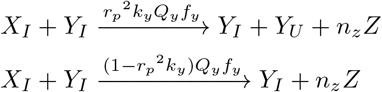

If the parasite is unsuccessful at infecting the infected predator, we apply the suppress of predator reproduction as before by parameter *r*_*e*_. Thus, in the following reaction where the predator reproduces, *r*_*e*_ refers to the cost of exposure in this encounter and *r*_*p*_ refers to the cost of previous infection.

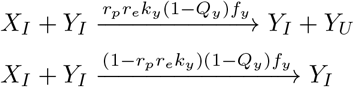

All individuals has an intrinsic death rate, with *d*_*x*_, *d*_*y*_ and *d*_*z*_ ∈ ℝ ^+^ denoting for the prey, predator and parasite respectively. While we already apply a cost of infection or exposure in reproduction, we assume the same intrinsic death for infected and uninfected individuals for the sake of simplicity.

However, this cost on death can be easily implemented if necessary.

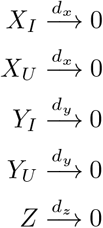

We use a standard Gillespie algorithm [34, 35] to perform a stochastic simulation of the time evolution of the prey and predator subtypes and the parasite. See Table 1 of the supplementary for the propensity equations and reactions.

**Table 1:** Parameter definitions.

| Parameter | Definition |
| --- | --- |
| $g_x$ | Reproduction rate of the prey |
| $r_x$ | Suppression of prey's reproduction due to parasite infection |
| $d_x$ | Intrinsic death rate of the prey |
| $K$ | Environmental carrying capacity of the prey |
| $S$ | Scaling factor for prey-parasite interactions |
| $n_z$ | Offspring per reproduction of the parasite |
| $d_z$ | Intrinsic death rate of the parasite |
| $f_y$ | Predation rate |
| $k_y$ | Reproduction rate of the predator |
| $r_p$ | Suppression of predator's reproduction due to parasite infection |
| $r_e$ | Suppression of predator's reproduction due to parasite exposure |
| $d_y$ | Intrinsic death rate of the predator |
| $Q_x$ | infection probability of parasite-prey |
| $Q_y$ | infection probability of parasite-predator |

### 2.2 Deterministic rate equations based on microscopic reactions

We can analyse the average population dynamics by writing down the deterministic equations based on the above reactions. We introduce *x, y* and *z* to represent the population abundance of the prey, predator and parasite respectively, where *x, y, z* ∈ ℝ ^+^. We use subscripts to distinguish the infected and susceptible prey and predator subtypes. We let *x*_*I*_ and *x*_*U*_ denote the infected and uninfected prey populations and *y*_*I*_ and *y*_*U*_ denote the infected and uninfected predator populations, and *x* and *y* for the total population. Thus, *x* = *x*_*I*_ + *x*_*U*_ and *y* = *y*_*I*_ + *y*_*U*_. We derive a set of ordinary differential equations from the microscopic reactions, where 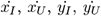, and *ż* denote the rates of change of corresponding populations with respect to time.

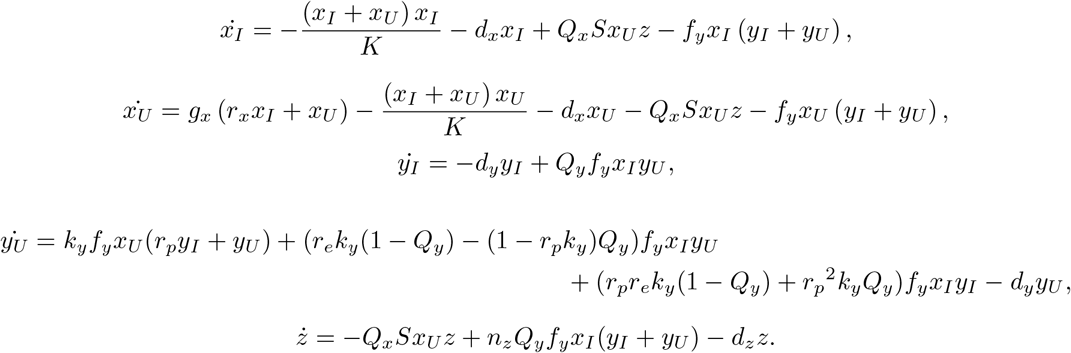

## 3 Results

### 3.1 Equilibrium analysis for the deterministic system

When the parasite enters the predator-prey system, to become endemic, it must have a basic reproduction value greater than unity. Given the parasite can only reproduce in an infected predator, (Figure 1), we show in section 2.1 of the supplementary that the infected predator population depends on the infected prey population and the infected prey population depends on the presence of the parasite, *z >* 0. We note here that a feature of the deterministic system considered is the existence of all species when *z >* 0.

**Figure 1.**
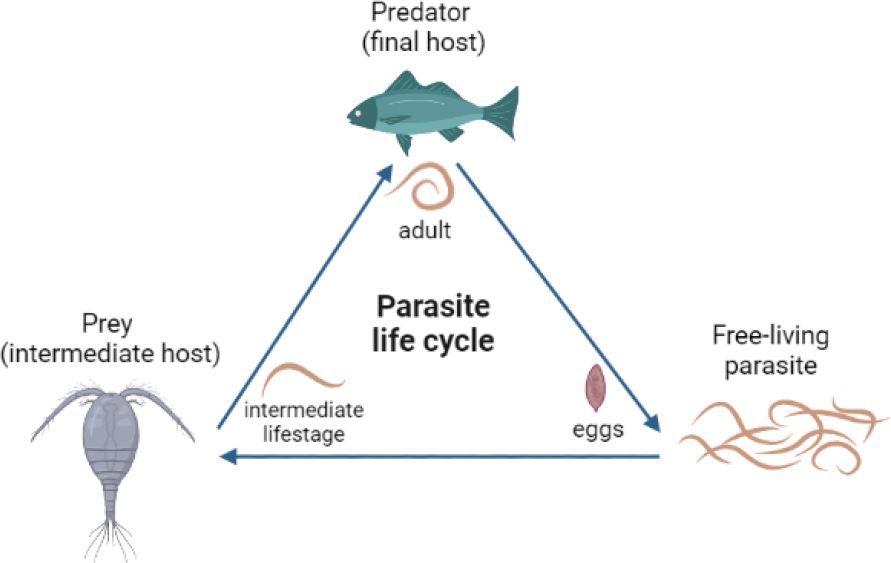
The species interactions in our model. The life cycle of a trophically transmitted parasite through two hosts under a predator-prey system.

When, *z >* 0, we have two scenarios, stable coexistence of all species or chaotic dynamics, as *t* → ∞. We can obtain the equilibria when 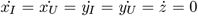 for *x*_*I*_, *x*_*U*_, *y*_*I*_, *y*_*U*_, *z >* 0. We show in section 2.2 of the supplementary the infected prey population steady state, *x*_*I*_^∗^, is a solution to the following polynomial.

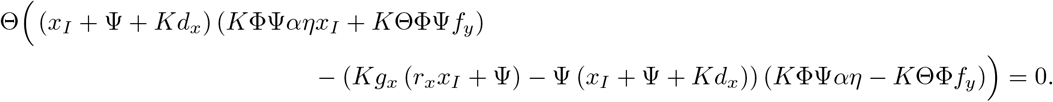

Here we have functions 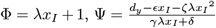 and Θ = *α*Ψ + *d*_z_, where *α* = *Q*_*x*_ *S, β* = *Q*_*y*_ *f*_*y*_, *γ* = *r*_*p*_*k*_*y*_*f*_*y*_, *δ* = *k*_*y*_*f*_*y*_, ϵ = (*r*_*e*_*k*_*y*_(1 − *Q*_*y*_) − (1 − *r*_*p*_*k*_*y*_)*Q*_*y*_)*f*_*y*_, *ζ* = (*r*_*p*_*r*_*e*_*k*_*y*_(1 − *Q*_*y*_) + *r*_*p*_^2^*k*_*y*_*Q*_*y*_)*f*_*y*_, *η* = *n*_*z*_*Q*_*y*_*f*_*y*_ and 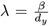. From *x*_*I*_^∗^, we determine the remaining steady state populations *x*_*U*_ ^∗^, *y*_*I*_^∗^, *y*_*U*_ ^∗^, *z*^∗^ as the following.

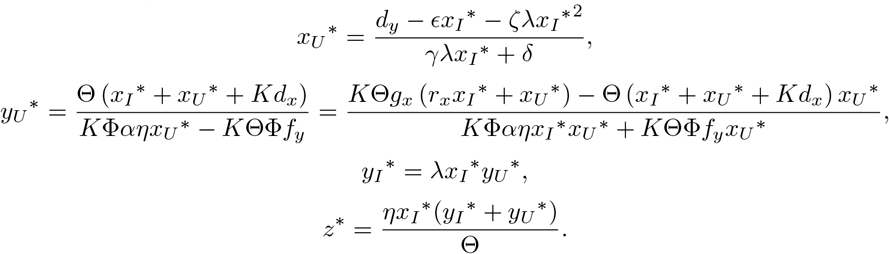

We observe a coexistence of all species, where our stochastic simulations fluctuate around the numerical solutions of the rate equations and converge to the analysed equilibrium when time increases (Figure 2). For reproduction suppression of infected predators, *r*_*p*_ = 0, the dynamics from the deterministic model are cyclic, as the predator population always exists. The stochastic simulations give different predictions: the predator goes extinct, followed by the parasite, and the prey population exists at carrying capacity in the absence of the predator (Figure S7).

**Figure 2.**
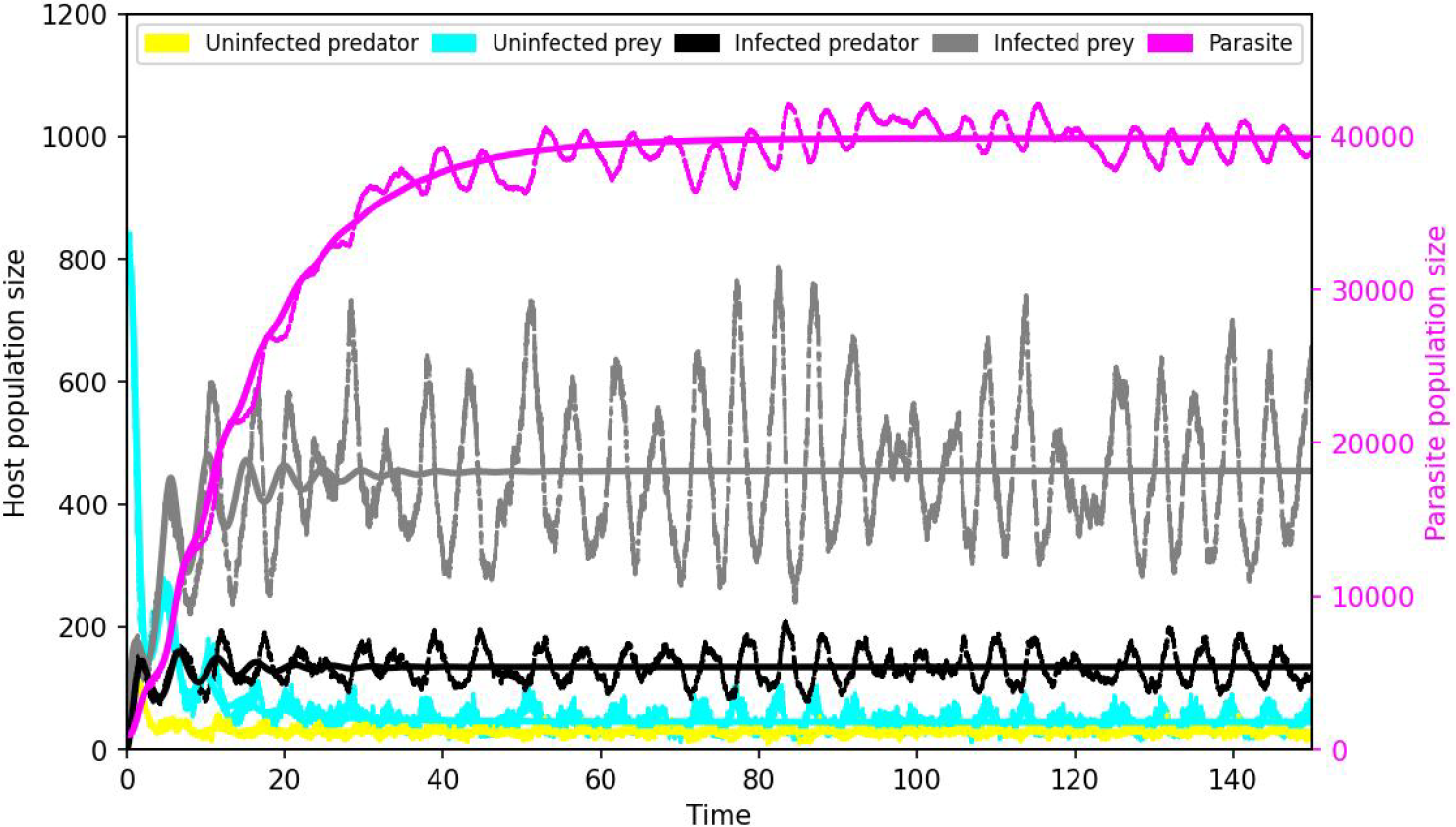
Population dynamics. The yellow colour shows the uninfected predator, the cyan colour shows the uninfected prey, the black colour shows the infected predator, the grey colour shows the infected prey, the magenta colour shows the parasites. The solid lines show the population dynamics according to deterministic simulations and the dashed lines according to stochastic simulations. Parameter values: *g*_*x*_ = 2, *r*_*x*_ = 1, *d*_*x*_ = 0.1, *K* = 2000, *S* = 0.0005, *n*_*z*_ = 6, *d*_*z*_ = 0.09, *f*_*y*_ = 0.01, *k*_*y*_ = 0.2, *d*_*y*_ = 1, *r*_*e*_ = 1, *r*_*p*_ = 1, *r*_*x*_ = 1, *Q*_*y*_ = 1 and *Q*_*x*_ = 1. Initial conditions: uninfected predator = 100, uninfected prey = 800, infected predator = 0, infected prey = 0 and parasite = 1000.

### 3.2 Coexistence of parasite, predator, and prey

In the absence of fitness costs, the coexistence of the three species is more likely with increasing infection probabilities of both host populations (Figure 3A, D). This is because parasites with complex life cycles must be efficient at infecting both trophic levels to reach maturity and reproduce. Interestingly, the three species coexist in a narrower range of infection probabilities of the predator host than of the prey host, with a threshold of *Q*_*y*_ *>* 0.25 and *Q*_*x*_ *>* 0.07, respectively. Given that the parasite depends on both hosts equally, these differences could stem from the predator population being smaller than the prey’s.

**Figure 3.**
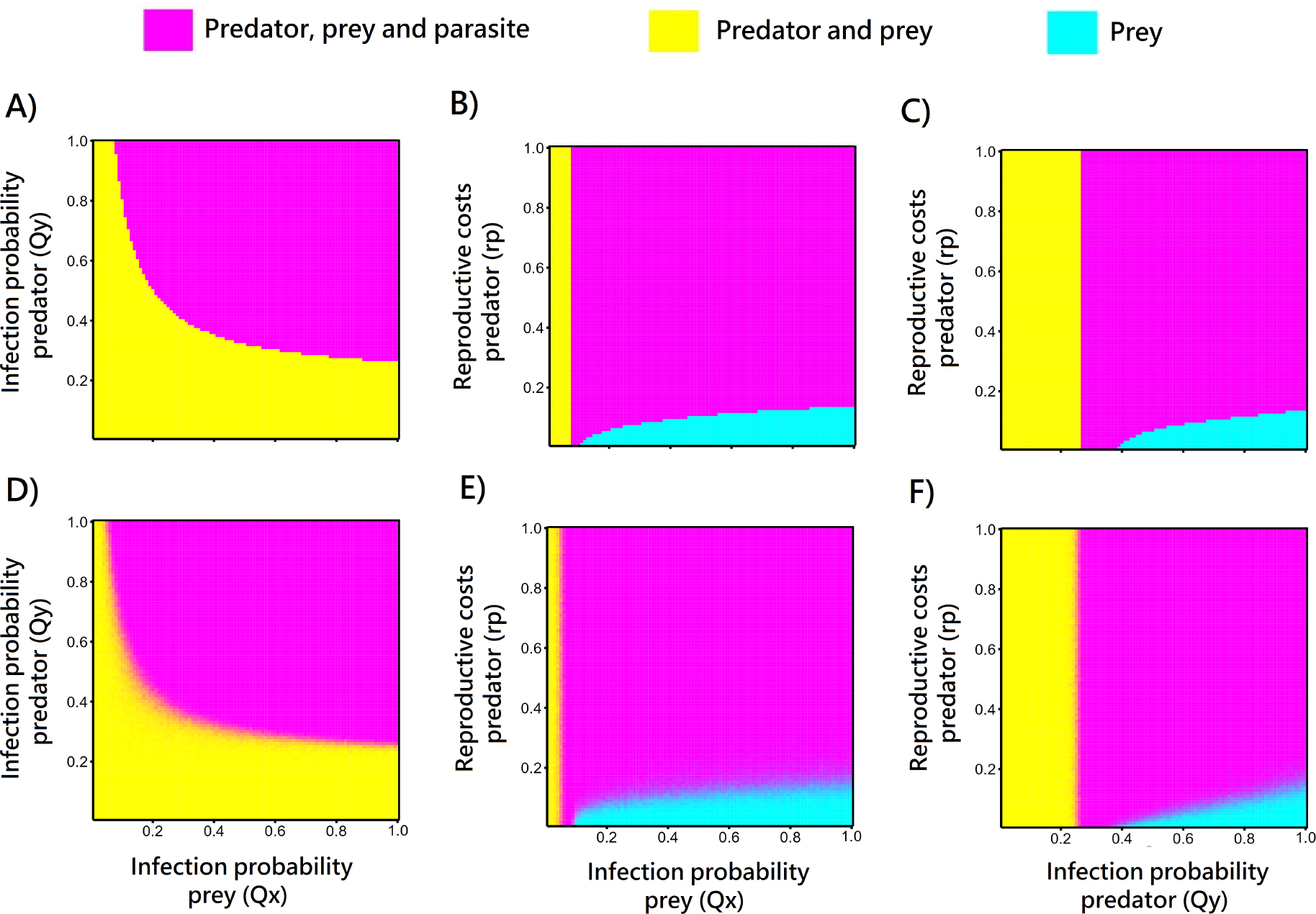
Patterns of species coexistence. The top panels show the deterministic simulations and the bottom panels show the stochastic simulations. In the stochastic simulations, each parameter combination displays the average value from 100 independent realisations over a time period of 130-150 time steps for each realisation. Parameter values: *g*_*x*_ = 2, *d*_*x*_ = 0.1, *K* = 2000, *S* = 0.0005, *n*_*z*_ = 6, *d*_*z*_ = 0.09, *f*_*y*_ = 0.01, *k*_*y*_ = 0.2, *d*_*y*_ = 1, *r*_*e*_ = 1, *r*_*x*_ = 1. Panels A and D show *r*_*p*_ = 1; Panels B and E show *Q*_*y*_ = 1; panels C and F show *Q*_*x*_ = 1. Initial conditions: uninfected predator = 100, uninfected prey = 800, infected predator = 0, infected prey = 0 and parasite = 1000. Note that *r*_*p*_ = 0 refers to high reproductive costs and *r*_*p*_ = 1 refers to the absence of reproductive costs.

When considering different degrees of fitness costs on the infected predator, we found that the stochastic simulations give different predictions than those from the deterministic simulations (Figure 3). For high reproductive costs on the predator, and abundant infected prey resulting from high values of *Q*_*x*_, the predator population was very low. We class that region of the parameter space as non-coexistence because the high suppression of the predator population results in the population being maintained at very low levels. At such low levels the predator population is vulnerable to extinction as observed in the stochastic system (Figure 3E, F). The dynamics of this parameter space are discussed in section 2.3 of the supplementary.

In the stochastic model, demographic fluctuations lead to the extinction of the predator when its population size drops because of high parasite virulence in both hosts (Figure S8A). Consequently, the parasite also undergoes extinction. With high reproductive costs on infected predators, the probability of predator-parasite coextinction increases with higher probabilities of the parasite infecting the prey, with a threshold of *Q*_*x*_ *>* 0.09 and *r*_*p*_ *<* 0.4 (Figure 3E). The same happens with the probabilities of the parasite infecting the predator, with a threshold of *Q*_*y*_ *>* 0.32 and *r*_*p*_ *<* 0.31 (Figure 3F). With high reproductive costs on infected prey, the probability of predator-parasite coextinction increases with higher infection probabilities of the prey, with a threshold of *Q*_*x*_ *>* 0.22, and *r*_*x*_ *<* 0.49 (Figure S8B). The same happens with higher infection probabilities of the predator, with a threshold of *Q*_*y*_ *>* 0.37 and *r*_*x*_ *<* 0.42 (Figure S8C). These results show the impact of the reproductive costs on one host scales up with the infection probability of the other host, i.e., an indirect parasite effect.

### 3.3 Frequency of uninfected and infected host subtypes

We consider the equilibrium points of infected and uninfected host subpopulations when the predator, prey, and parasite coexist (Figure 4). As expected, infected individuals are more frequent than their uninfected counterparts with increasing infection probabilities and a threshold of *Q*_*x*_ *>* 0.18 and *Q*_*y*_ *>* 0.36, in both the deterministic and stochastic simulations (Figure 4A, D). Similarly, uninfected hosts coexist with the parasite at higher frequencies than their infected counterparts with low infection probabilities. Interestingly, we found that all combinations of subpopulations’ coexistence are possible, albeit at very different likelihoods. For instance, the most common outcome predicted shows a higher number of infected predators combined with a higher amount of uninfected prey. This combination of host frequencies happens when the parasite is more efficient at infecting the predator than the prey, with a threshold of *Q*_*x*_ = [0.12 − 0.69] and *Q*_*y*_ *>* 0.41 (Figure 4A, D). In the absence of reproductive costs on infected predators, this outcome happens in a small parameter space of infection probabilities of the prey *Q*_*x*_ = [0.12− 0.18] (Figure 4B, E). Contrarily, with high reproductive costs on infected predators, this parameter space increases, and the infection probabilities of both hosts become less relevant (Figure 4B, C, E, F). This suggests that the prey can recover from infection when parasite pressure leads to few predators, as it relaxes the predation pressures on the prey. In addition, when the predator is less efficient at consuming the prey, the predator population is unable to recover from infection, regardless of the type of prey that is available.

**Figure 4.**
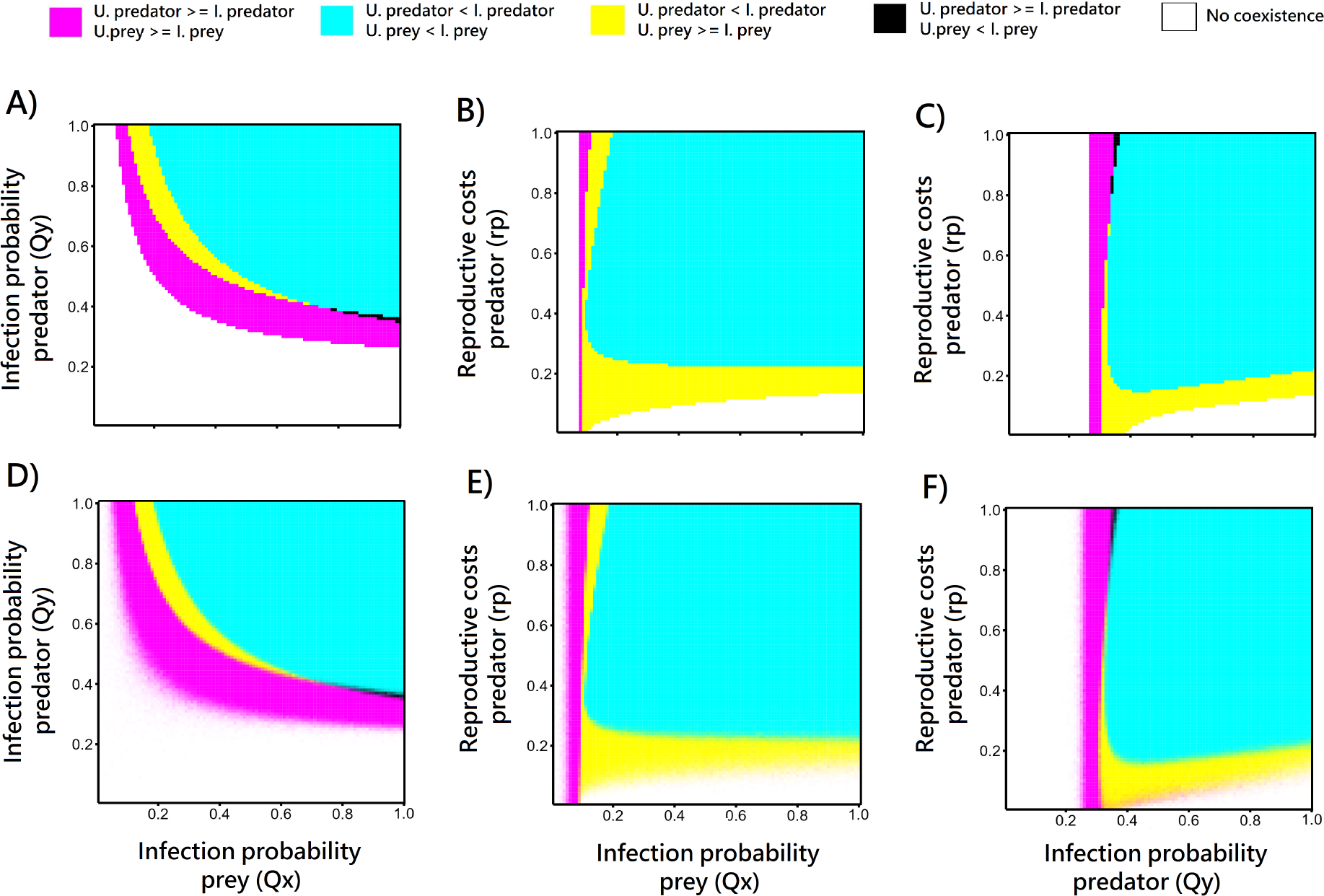
Comparing the relative frequencies of infected and uninfected prey and predators. The top panels show the deterministic simulations and the bottom panels show the stochastic simulations. In the stochastic simulations, each parameter combination displays the average value from 100 independent realisations over a time period of 130-150 time steps for each realisation. Parameter values: *g*_*x*_ = 2, *d*_*x*_ = 0.1, *K* = 2000, *S* = 0.0005, *n*_*z*_ = 6, *d*_*z*_ = 0.09, *f*_*y*_ = 0.01, *k*_*y*_ = 0.2, *d*_*y*_ = 1, *r*_*e*_ = 1, *r*_*x*_ = 1. Panels A and D show *r*_*p*_ = 1; Panels B and E show *Q*_*y*_ = 1; panels C and F show *Q*_*x*_ = 1. Initial conditions: uninfected predator = 100, uninfected prey = 800, infected predator = 0, infected prey = 0 and parasite = 1000. Note that *r*_*p*_ = 0 refers to high reproductive costs and *r*_*p*_ = 1 refers to the absence of reproductive costs.

A high frequency of infected prey combined with a high frequency of uninfected predators is the least common outcome, yet an interesting one (Figure 4A, C, D, F). This combination of host frequencies occurs in a parameter space in which the parasite is less efficient at infecting the predator than the prey, combined with low reproductive costs on infected predators, with a threshold of *Q*_*y*_ = 0.33 − 0.38, and *r*_*p*_ *>* 0.6, respectively. Due to the complex life cycle of the parasite, the predator must eat infected prey to be exposed to the parasite. When there is a high availability of infected prey, the transmission of the parasite to the predator ultimately depends on the predator’s resistance/susceptibility, ultimately representing their immunity. When the parasite’s virulence is low and the prey is not often infected, the parasite will not reach the predator, independently of the predator’s immunity. This explains why we do not see the coexistence of infected prey and uninfected predators at high frequencies according to any probability of the parasite infecting the prey. In addition, with low reproductive costs on infected predators, the predator population recovers rapidly from infection because we assume that the parasite is not vertically transmitted to the next generation.

To investigate further the conditions leading to a high frequency of infected prey and uninfected predator, we tested the effects of the parameters when *Q*_*y*_ = 0.35, which is the range of infection probability of the predator leading to these frequencies of host subpopulations. We found that the variation in the probability of prey infection and reproductive costs on the prey become practically irrelevant and that this outcome is mainly driven by a low probability of predator infection combined with low reproductive costs on the predator (Figure 5B). This interesting result suggests that the probability of the parasite infecting the prey is ultimately mediated by predation pressures which are, in turn, mediated by the parasite-mediated selection, hence revealing an indirect parasite effect.

**Figure 5.**
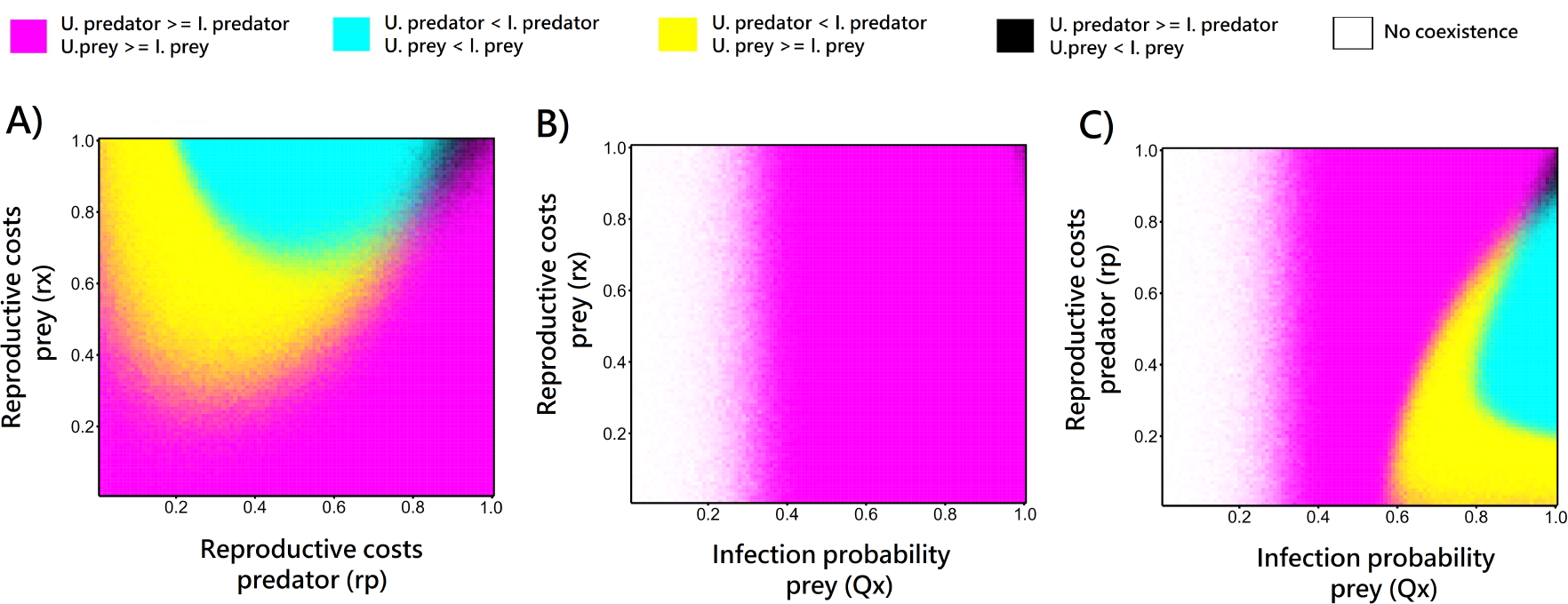
Comparing the relative frequencies of infected and uninfected prey and predators. The panels show stochastic simulations. Each parameter combination displays the average value from 100 independent realisations over a time period of 130-150 time steps for each realisation. Parameter values: *g*_*x*_ = 2, *d*_*x*_ = 0.1, *K* = 2000, *S* = 0.0005, *n*_*z*_ = 6, *d*_*z*_ = 0.09, *f*_*y*_ = 0.01, *k*_*y*_ = 0.2, *d*_*y*_ = 1, *r*_*e*_ = 1, *Q*_*y*_ = 0.35. Panel A shows *Q*_*x*_ = 1; Panel B shows *r*_*p*_ = 1; panel C shows *r*_*x*_ = 1. Initial conditions: uninfected predator = 100, uninfected prey = 800, infected predator = 0, infected prey = 0 and parasite = 1000. Note that *r*_*p*_ = 0 (and *r*_*x*_ = 0) refers to high reproductive costs and *r*_*p*_ = 1 (and *r*_*x*_ = 1) refers to the absence of reproductive costs.

## 4 Discussion

Growing evidence suggests that parasites mediate trophic interactions and that predators regulate hostparasite interactions [25, 26]. Yet, the interplay of those dynamics remains understudied, particularly when parasites have complex life cycles [13]. To address this knowledge gap, we designed an individual-based model of a complex predator-prey-parasite system, where the parasite is transmitted trophically from infected prey to predators. Using simulations, we show that the infection probabilities of both host species and parasite virulence (reproductive costs on infected hosts) determine the coexistence of the three species. We compared our stochastic simulation results with the mathematical analysis of deterministic rate equations. In almost all parameter regimes, our stochastic simulations agree with deterministic predictions well. In the border regions (e.g., between coexistence and extinction in the parameter space), while only one outcome is possible under deterministic dynamics for each parameter set, both coexistence and extinction can happen in different stochastic realisations. This demonstrates the relevance of demographic fluctuations in species coexistence. When decomposing the host populations into uninfected and infected subpopulations, we found multiple stable states of the three species which are determined by the infection probabilities and parasite virulence across the trophic levels.

Generally, we show that the three species are more likely to coexist with increasing infection probabilities of both hosts because parasites of complex life cycles need to infect both trophic levels to develop and reproduce [36, 37]. Interestingly, our results further show that, for the persistence of the parasite, the probability of infecting the predator is more critical than the probability of infecting the prey. Given that the parasite depends on both hosts equally, these differences could stem from the different population sizes of the predator and prey. We suggest that the parasite needs to be more efficient at infecting the predator because the encounter probabilities between uninfected predators and infected prey are lower than those between free-living parasites and uninfected prey. An alternative, non-excluding, explanation is that the availability of infected or uninfected prey becomes less relevant when the parasite is inefficient at infecting the predator, i.e., with low probabilities of infecting the predator.

Previous deterministic models of parasite dynamics within simple communities predict that parasites should generally go extinct before their hosts [38]. Here, using complex communities, we show similar patterns whereby, in most cases, the parasite goes extinct before the predator, but the prey always survives. Stochastic simulations however reveal that the parasite alone, or both the predator and the parasite, can go extinct when the fitness costs of infection are high on the predator combined with medium to high probabilities of the parasite infecting both hosts. This result is consistent with theoretical work on complex predator-prey-parasite systems (where the parasite infects only the prey), showing that when the prey population is infected at a low rate, and the predator has low reproductive potential, the extinction of both the predator and the parasite can happen [39]. We suggest, however, that the underlying mechanisms are different when considering a parasite that mandatorily infects both trophic levels. In our simulations, we show that indeed predator-parasite coextinctions happen with low reproductive potential of the predator, however, in combination with medium to high probabilities of the parasite infecting both hosts. This pattern emerges because, in that parameter space, the predator population size is low. Furthermore, our results indicate that, due to the parasite’s complex life cycle, the fitness of one host (estimated in the form of reproductive success) can change with the probability of the other host getting infected. These complex interactions are reflected in our stochastic simulations, where the likelihood of predator-parasite coextinction increases with high infection probabilities of the predator combined with high fitness costs on infected prey, as well as with high infection probabilities of the prey combined with high fitness costs on infected predators. These results illustrate an indirect parasite effect on predator-prey dynamics and highlight the ecological relevance of complex species interactions.

To better understand the dynamics of the parasite’s demography, we decoupled the predator and prey populations into uninfected and infected subpopulations when the parasite coexists with the predator and the prey at a stable equilibrium. The most common outcome of our simulations reveals a high frequency of infected predators coexisting with a high frequency of infected prey. As expected, the frequencies of infected hosts are higher with increasing infection probabilities and are lower with decreasing infection probabilities. Intuitively, due to the parasite’s complex life cycle, we expected the frequencies of infected and uninfected predator subpopulations to match with those of the prey. Yet, although less common, in some parameter spaces, this is not the case. Particularly, with high reproductive costs on infected predators, the predator population is less likely to recover from infection. In that case, the infection probabilities of both hosts become less relevant in the frequency of infected predators (Figure 4B), illustrating a direct parasite effect on the predator. Also, as the frequency of infected predators increases, their population size drops, hence relaxing the predation pressure on the prey, illustrating an indirect parasite effect on the prey through the predator. Consequently, the prey population size recovers mostly with uninfected prey. Together, the intersection of parasite-mediated and predation-mediated selection, explains the higher number of uninfected prey and infected predator individuals.

We found that the least likely combination of host subpopulations is a high frequency of infected prey coexisting with a high frequency of uninfected predators. While this outcome is interesting, it is rare and happens in a narrow parameter space of infection probability of the predator *Q*_*y*_ = [0.33−0.38], combined with low reproductive costs on infected predators *r*_*p*_ *>* 0.6. Given that our model does not assume vertical transmission of the parasite to the prey or predator offspring, we expected the prey population to recover from infection in the absence of reproductive fitness costs. Surprisingly, the likelihood of this outcome increases with low reproductive costs on infected prey. We suggest that the prey recovers slower from infection due to strong predation pressures in this parameter space, particularly as the infection probability is higher on the prey than on the predator. In addition, we found that in the absence of reproductive costs on infected predators, the combination of high frequencies of infected prey and uninfected predators does not happen in any parameter space of infection probabilities or reproductive costs on the prey population (Figure 5B). This illustrates the role of the predator’s demography in mediating the prevalence of infection in the prey. This result is coherent with the healthy herd hypothesis, which proposes that the dynamics of prey-parasite interactions are determined by the predator population size and parasite virulence [40]. Also, experimental evidence shows that the parasite’s impact on the population size of an intermediate consumer is obscured by that of the predator when predators are abundant [39]. which is a characteristic we found often in our system.

In natural systems, parasites can be virulent, and yet sometimes do not cause significant changes in host density [41]. For instance, Duffy and Hall (2008) theoretically tested the effects of two parasites, the bacterium *Spirobacillus cienkowskii* and the yeast *Metschnikowia bicuspidate*, on the common freshwater invertebrate *Daphnia denifera* [41]. While both parasites are known to be virulent, only bacterial epidemics led to significant changes in the host density. Interestingly, they found that the rapid evolution of host resistance to the yeast parasite combined with predation mostly on infected hosts decreased the prevalence of infection and minimised host density decline [41]. Further theoretical work on eco-epidemiological models, where the parasite infects the prey population, showed that the predators’ preference for uninfected and infected prey individuals plays a significant role in the system dynamics and stability [42, 43]. We suggest that these dynamics may be different in systems where the parasite is transmitted trophically because the infection probabilities and parasite virulence can vary across trophic levels [44].

Based on our results, we propose that the combination of host subpopulations in different trophic levels may have important ecological consequences. For instance, if a predator evolves the capacity to eat preferentially uninfected prey, high predation pressures will result in a prey population of highly abundant infected individuals [40]. The resulting high incidence of infected prey may lead to a higher likelihood of transmission of the parasite to other predators that lack a discriminative feeding behaviour. We suggest that the transmission of these parasites to other predator species will ultimately depend on multiple factors such as parasite virulence and the immunity of those hosts. Similarly, a high incidence of infected predators or a high incidence of infected prey may lead to trophic cascades [24] with potentially important consequences at the community level [25, 26].

Overall, we demonstrate that the interplay of direct and indirect parasite effects along a food web is a common driver of the prevalence of infections. The parasite can impact the host population directly through the reproductive costs resulting from infection, or indirectly via changing the demography of the interacting species, i.e., the predator or the prey of the host. The combination of infection probabilities and reproductive costs on infected hosts of different trophic levels determines the internal stability of the system because parasites change the demography of the host populations (i.e., a direct parasite effect) and, consequently, predator-prey dynamics (i.e., an indirect parasite effect). Moreover, by analysing the dynamics of different host subpopulations, we provide quantitative data on the prominent features of a multi-species system driving the dynamics of a trophically transmitted parasite. In some parameter spaces, we show that both extinction and coexistence can happen in different stochastic realisations. Although we focus on a static predator-prey-parasite system, the stochastic uncertainties might lead to completely different evolutionary outcomes when mutations and coevolution happen. We propose that integrating the evolution of host resistance and parasite infectivity in complex models may reveal important aspects of these complex interactions because the rates of infection vary according to host-parasite coevolution. Particularly, hosts from different trophic levels may coevolve with the parasite differently, according to their population sizes and parasite virulence. Yet, our study provides new insights into the mechanisms driving the dynamics of trophically transmitted parasites, with important implications for forecasting and managing the spread of diseases in natural ecosystems.

## Supplementary Material

### 1 List of reactions in our stochastic simulations

We implement the stochastic dynamics of our system with Gillespie algorithm. The propensity functions and the reactions required described in section 2.1 of the main text are listed in table 1.

**Table 1:**
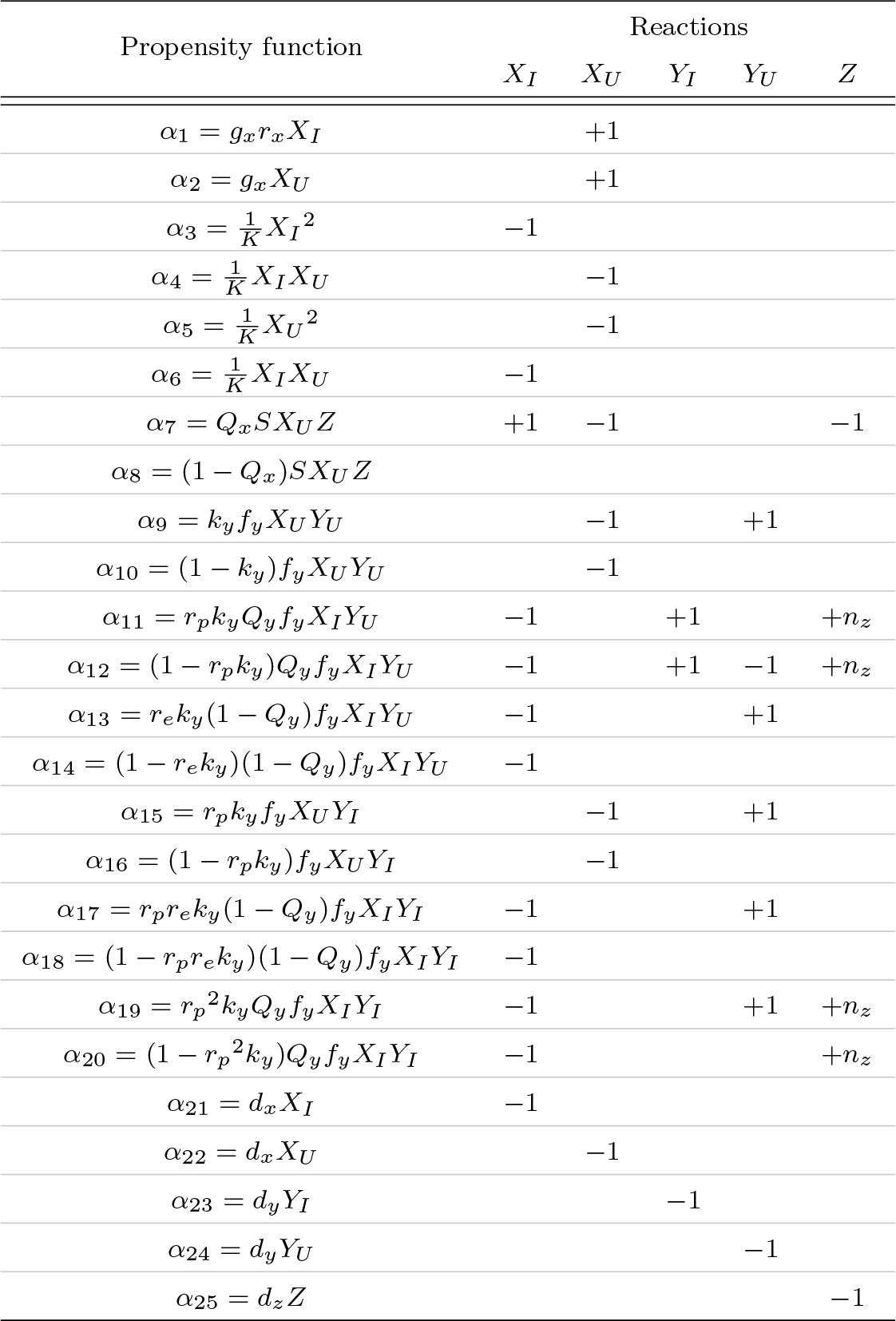
Propensity functions and reactions for stochastic simulations for predator-prey interactions with parasite present.

### 2 Deterministic analysis

As shown in the section 2.2 of the main text, we have the following set of equations for the average population dynamics

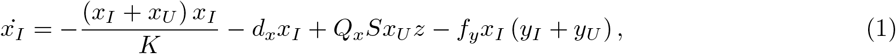

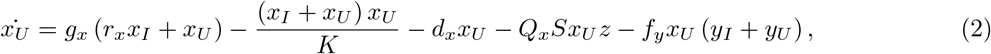

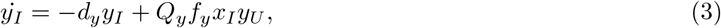

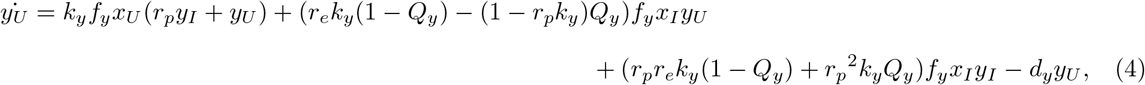

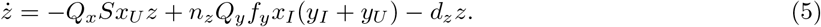

#### 2.1 The critical condition for the reproduction of the parasite

Since the parasite reproduces in the definitive host, the predator, the parasite can invade in the multiple host system and will be endemic when 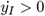 close to *t* = 0. We get:

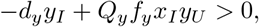

which we write as:

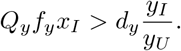

Initially the infected predator population is zero and the entire predator population is susceptible, so we have:

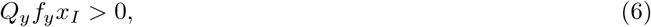

and we see that 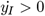 is dependant on *x*_*I*_ *>* 0. With a complex life cycle a parasite must first be sustained in the immediate host population to then become endemic in the definitive host population. Let us now consider 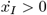 close to *t* = 0. From 1 we get:

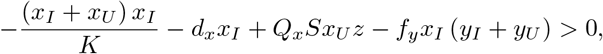

which becomes:

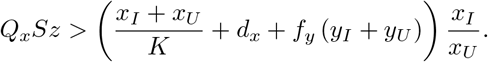

The infected prey population is zero at *t* = 0, so we get:

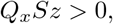

which in the presence of a parasite, *z >* 0, along with probability *Q*_*x*_ and *S >* 0 shows the parasite is sustained within the prey population since if 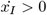 close to *t* = 0 then *x*_*I*_ *>* 0 close to *t* = 0. Now together with probability *Q*_*y*_ and *f*_*y*_ *>* 0, we conclude from 6, the parasite is always endemic within the predator population.

#### 2.2 Deriving the equilibria of our deterministic equations

We now determine the fixed points of the system described by 1 - 5. That is we determine when:

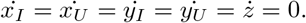

Again we are interested in when all species coexist that is when *x*_*I*_, *x*_*U*_, *y*_*I*_, *y*_*U*_, *z >* 0. Here we let *α* = *Q*_*x*_*S, β* = *Q*_*y*_*f*_*y*_, *γ* = *r*_*p*_*k*_*y*_*f*_*y*_, *δ* = *k*_*y*_*f*_*y*_, *ϵ* = (*r*_*e*_*k*_*y*_(1 − *Q*_*y*_) − (1 − *r*_*p*_*k*_*y*_)*Q*_*y*_)*f*_*y*_, *ζ* = (*r*_*p*_*r*_*e*_*k*_*y*_(1 − *Q*_*y*_) + *r*_*p*_^2^*k*_*y*_*Q*_*y*_)*f*_*y*_ and *η* = *n*_*z*_*Q*_*y*_*f*_*y*_. Then we write the system described by 1 - 5 as:

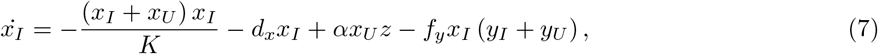

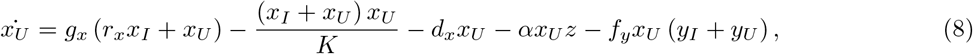

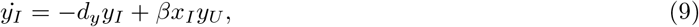

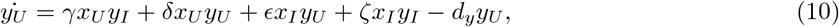

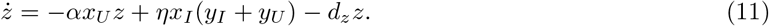

First let us consider 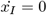, from 7 we get:

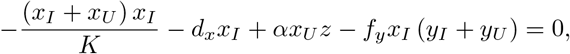

which we write as

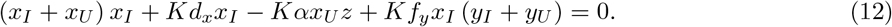

Next let us consider 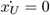, 8 yields:

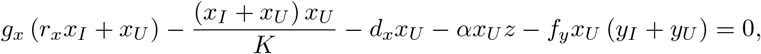

and we write:

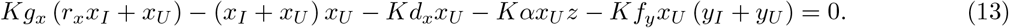

Now we consider *ż* = 0. From 11 we get:

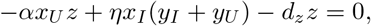

which we write as:

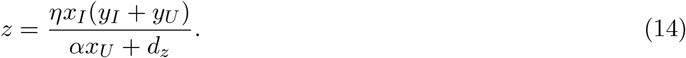

Let us introduce Θ a function of *x*_*U*_ where, Θ = *αx*_*U*_ + *d*_*z*_. Then 14 becomes:

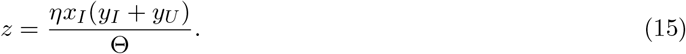

Now we substitute 15 into 12, which yields:

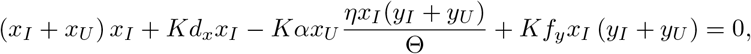

that we write as:

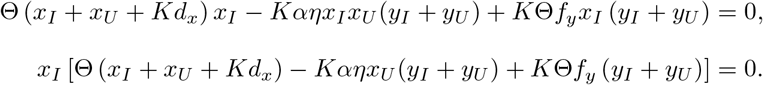

So we have the trivial solution *x*_*I*_ = 0, which we ignore, or the solution:

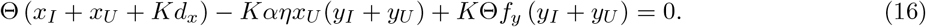

We next substitute 15 into 13 and we get:

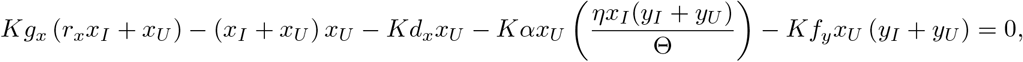

that we write as:

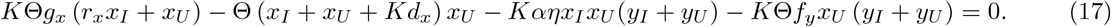

Now let us consider when 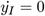, from 9 we get:

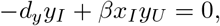

which we write as:

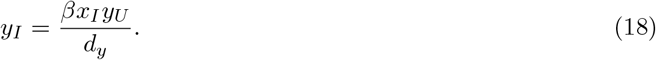

Here we let 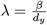 and we write 18 as:

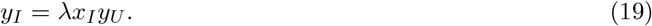

Now we substitute 19 into 16, which yields:

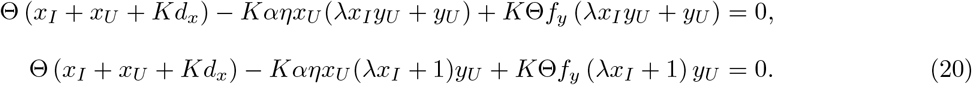

Here we introduce Φ a function of *x*_*I*_ where Φ = *λx*_*I*_ + 1. Now we write 20 as:

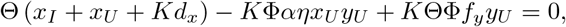

and finally as:

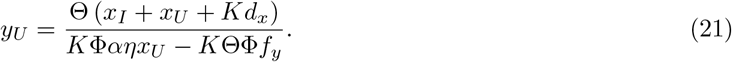

Next we substitute 19 into 17. Here we get:

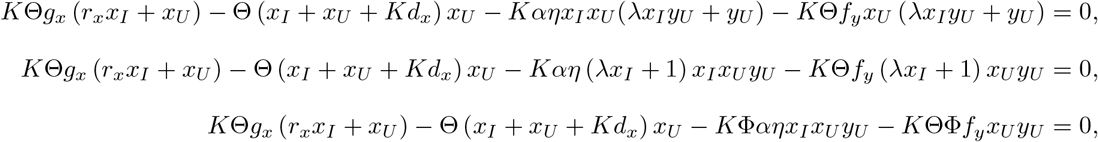

that we write as:

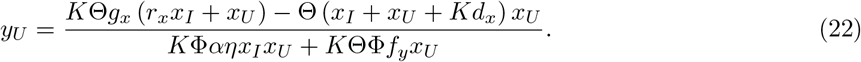

Now we substitute 21 into 22, which yields:

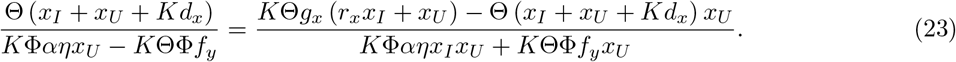

We write 23 as:

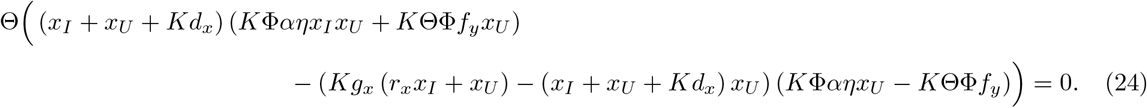

Finally let us consider when 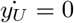, from 10 we get:

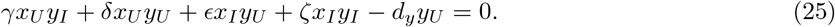

Now substitute 19 into 25, which yields:

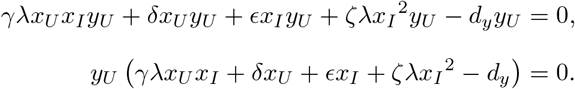

Here we have the trivial solution *y*_*U*_ = 0, which we again ignore, or the solution:

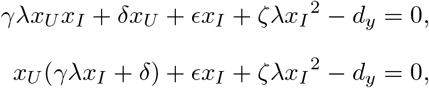

which we write as:

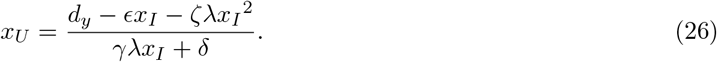

Let 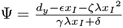 be a function of *x*_*I*_. Then we can write Θ as a function of *x*_*I*_ where:

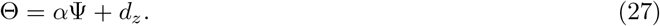

Now when we substitute 26 into 24 we get a function in terms of *x*_*I*_ as follows:

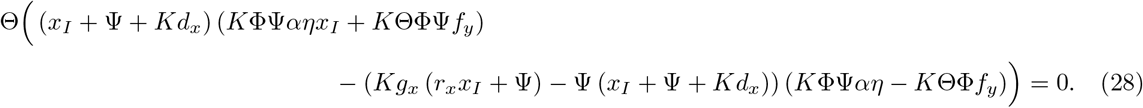

We can solve 28 numerically and visual inspection of these dynamics can be used to determine the steady state population for the infected prey, which we denote *x*_*I*_^∗^. We can then determine the remaining equilibria values using 26 for the susceptible prey, 21 or 22 for the susceptible predator, 19 for the infected predator and 15 for the parasite, which we denote *x*_*U*_ ^∗^,*y*_*U*_ ^∗^,*y*_*I*_^∗^ and *z*^∗^ respectively. Now we have the fixed point (*x*_*I*_^∗^, *x*_*U*_ ^∗^, *y*_*I*_^∗^, *y*_*U*_ ^∗^, *z*^∗^) which corresponds to the asymptotic populations where there is coexistence of all species.

### 2.3 Coexistence of all three species in deterministic dynamics

**Figure 1:**
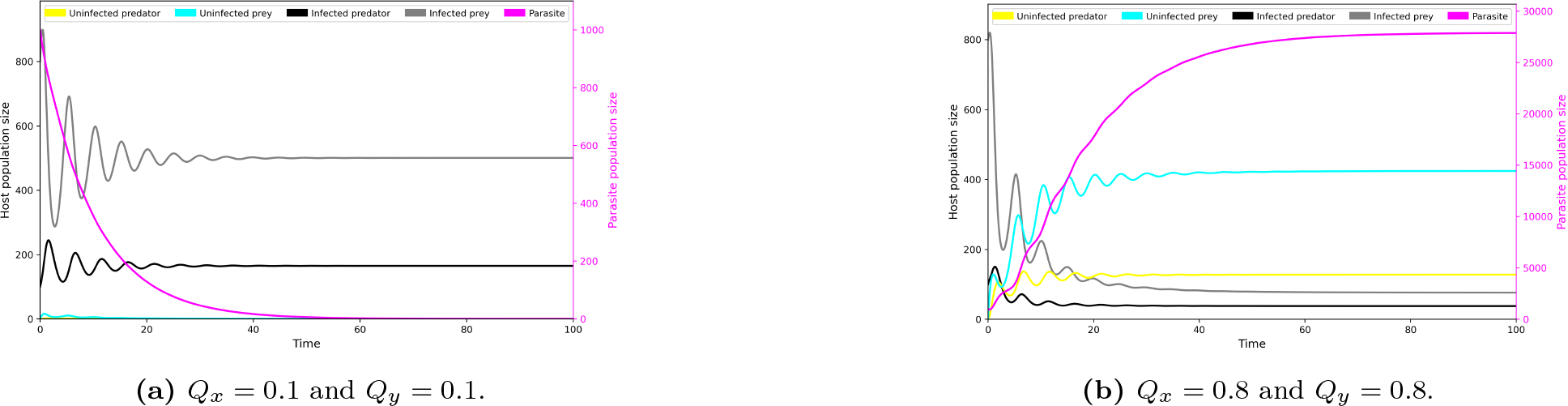
Comparing the behaviour of the system, according to numerical approximation of deterministic equations. Parameter values: *g*_*x*_ = 2, *d*_*x*_ = 0.1, *K* = 2000, *S* = 0.0005, *n*_*z*_ = 6, *d*_*z*_ = 0.09, *f*_*y*_ = 0.01, *k*_*y*_ = 0.2, *d*_*y*_ = 1, *r*_*x*_ = 1, *r*_*p*_ = 1 and *r*_*e*_ = 1. Initial conditions: uninfected predator = 100, uninfected prey = 800, infected predator = 0, infected prey = 0 and parasite = 1000.

We consider parameters *Q*_*x*_, *Q*_*y*_ and *r*_*p*_, the probability that the parasite infects the prey, the probability that the parastite infects the predator and the suppression the parasite has on the infected predators ability to reproduce.

**Figure 2:**
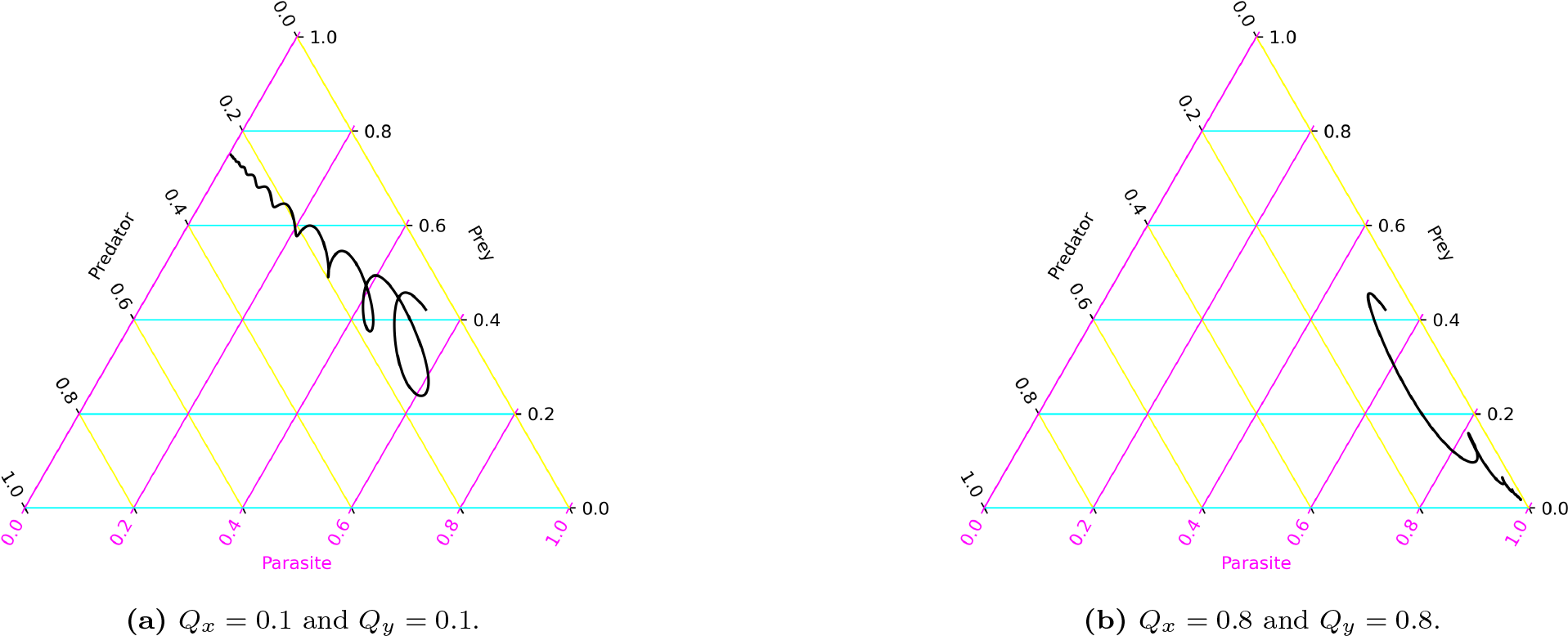
Simplex of species, according to numerical approximation of deterministic equations. Parameter values: *g*_*x*_ = 2, *d*_*x*_ = 0.1, *K* = 2000, *S* = 0.0005, *n*_*z*_ = 6, *d*_*z*_ = 0.09, *f*_*y*_ = 0.01, *k*_*y*_ = 0.2, *d*_*y*_ = 1, *r*_*x*_ = 1, *r*_*e*_ = 1 and *r*_*p*_ = 0.1. Initial conditions: uninfected predator = 100, uninfected prey = 800, infected predator = 0, infected prey = 0 and parasite = 1000.

For low values of *Q*_*x*_ and *Q*_*y*_ the parasite is not able to sustain itself in the system and we have extinction of the parasite and infected prey and predator populations. The system is then reduced to a damped Lotka-Volterra system, see Figure S1(a).

**Figure 3:**
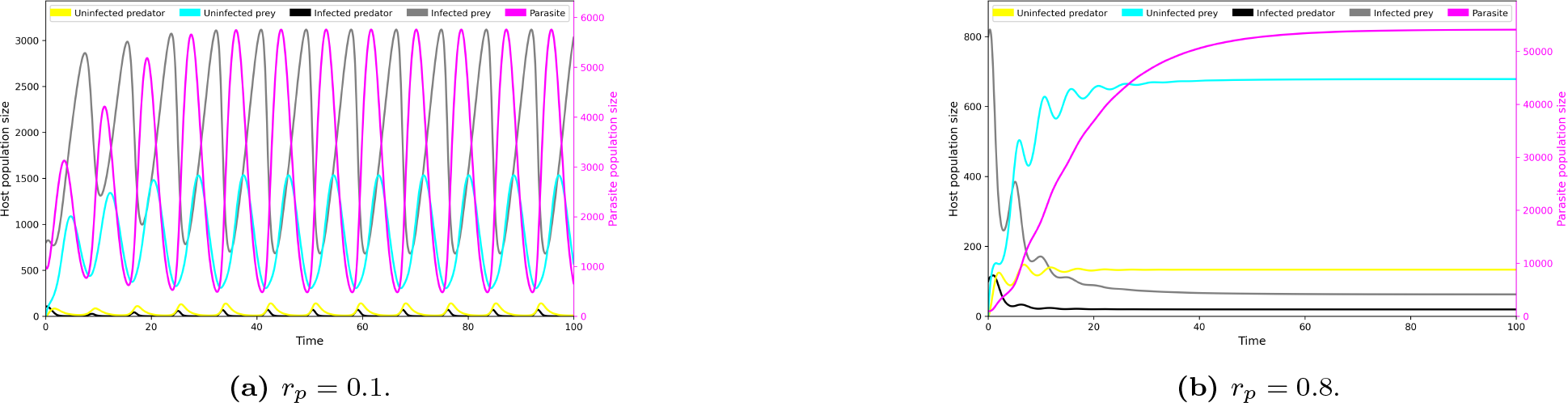
Comparing the behaviour of the system, according to numerical approximation of deterministic equations. Parameter values: *g*_*x*_ = 2, *d*_*x*_ = 0.1, *K* = 2000, *S* = 0.0005, *n*_*z*_ = 6, *d*_*z*_ = 0.09, *f*_*y*_ = 0.01, *k*_*y*_ = 0.2, *d*_*y*_ = 1, *r*_*x*_ = 1, *r*_*e*_ = 1, *Q*_*x*_ = 0.8 and *Q*_*y*_ = 1. Initial conditions: uninfected predator = 100, uninfected prey = 800, infected predator = 0, infected prey = 0 and parasite = 1000.

Figure S1(b) shows that for high values of *Q*_*x*_ and *Q*_*y*_ we have coexistence of all populations and the populations as *t* → ∞ converge to the equilbria values evaluated in the previous section. Figure S2 shows the trajectories for the cases described above.

**Figure 4:**
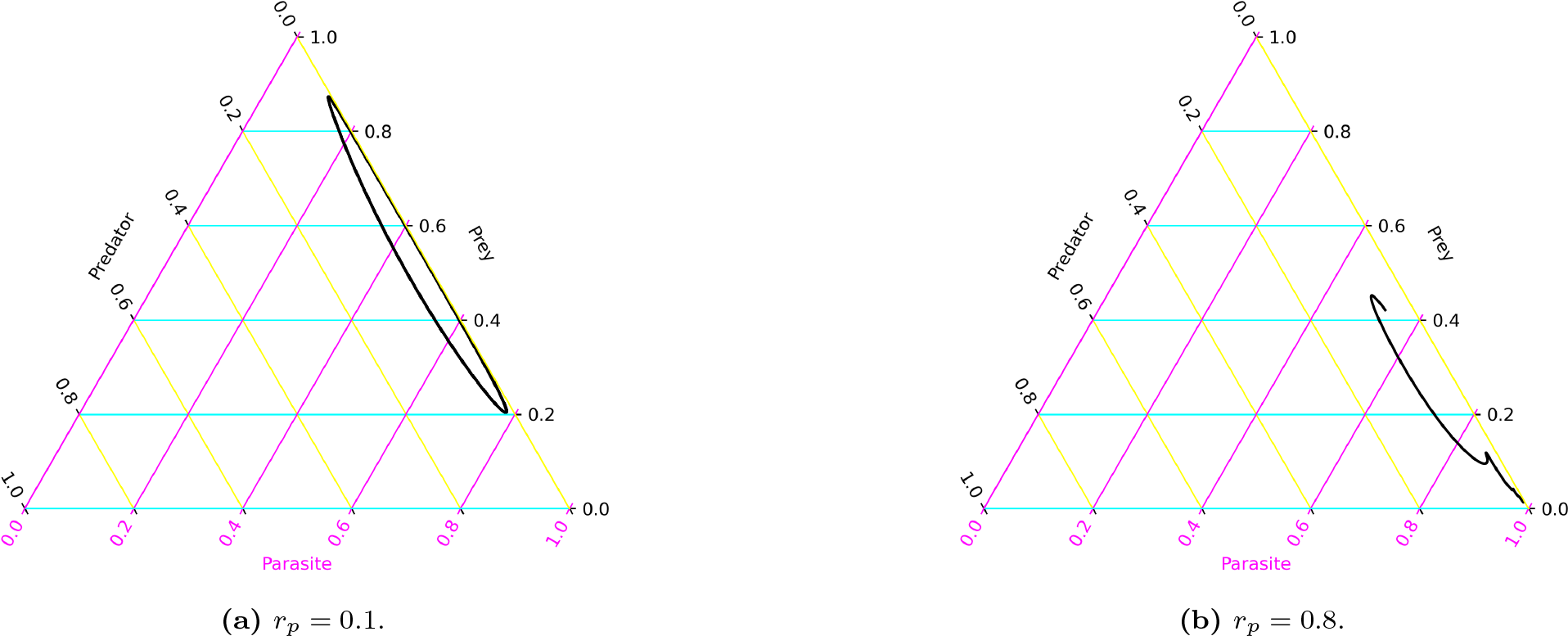
Simplex of species, according to numerical approximation of deterministic equations. Parameter values: *g*_*x*_ = 2, *d*_*x*_ = 0.1, *K* = 2000, *S* = 0.0005, *n*_*z*_ = 6, *d*_*z*_ = 0.09, *f*_*y*_ = 0.01, *k*_*y*_ = 0.2, *d*_*y*_ = 1, *r*_*x*_ = 1, *r*_*e*_ = 1, *Q*_*x*_ = 0.8 and *Q*_*y*_ = 1. Initial conditions: uninfected predator = 100, uninfected prey = 800, infected predator = 0, infected prey = 0 and parasite = 1000.

Next we consider *Q*_*x*_ and *r*_*p*_. Recall for low values of *Q*_*x*_ the parasite is not able to sustain itself in the system. For high *Q*_*x*_ and low *r*_*p*_ we have high suppression of the infected predator populations ability to reproduce and this results in low overall predator populations as with high *Q*_*x*_ the parasite has high virulence. The resulting behaviour in the system is cyclic as the predator population is reduced the prey population increases due to a decrease in predation, this in turn provides an abundant food supply for the predator and also for the parasite to infect the intermediate host. In the abundance of food, with significant infected subtype population, the predator population starts to increase until the suppression of the infected predators ability to reproduce drives a decrease in the predator population. Figure S4(a) shows the cyclic dynamics for high *Q*_*x*_ and low *r*_*p*_.

**Figure 5:**
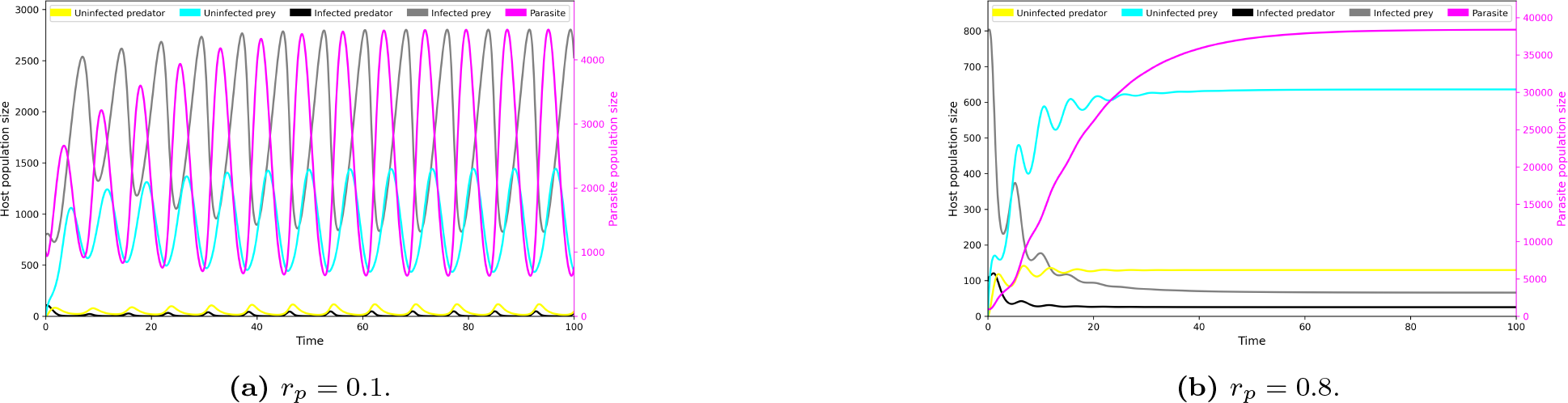
Comparing the behaviour of the system, according to numerical approximation of deterministic equations. Parameter values: *g*_*x*_ = 2, *d*_*x*_ = 0.1, *K* = 2000, *S* = 0.0005, *n*_*z*_ = 6, *d*_*z*_ = 0.09, *f*_*y*_ = 0.01, *k*_*y*_ = 0.2, *d*_*y*_ = 1, *r*_*x*_ = 1, *r*_*e*_ = 1, *Q*_*x*_ = 0.8 and *Q*_*y*_ = 1. Initial conditions: uninfected predator = 100, uninfected prey = 800, infected predator = 0, infected prey = 0 and parasite = 1000.

When we consider the system for high *Q*_*x*_ and high *r*_*p*_, that is low suppression of the infected predators ability to reproduce, we see coexistence of all populations and the populations as *t* → ∞ converge to the equilbria values evaluated in the previous section, see Figure S3(b). We observe similar dynamics to *Q*_*x*_ and *r*_*p*_ when we consider *Q*_*y*_ and *r*_*p*_ (Figure S5). Figure S6 shows the simplices of species populations for high *Q*_*y*_ with low and high *r*_*p*_ respectively. In (a) we observe cycles and in (b) we observe convergence to the fixed point.

**Figure 6:**
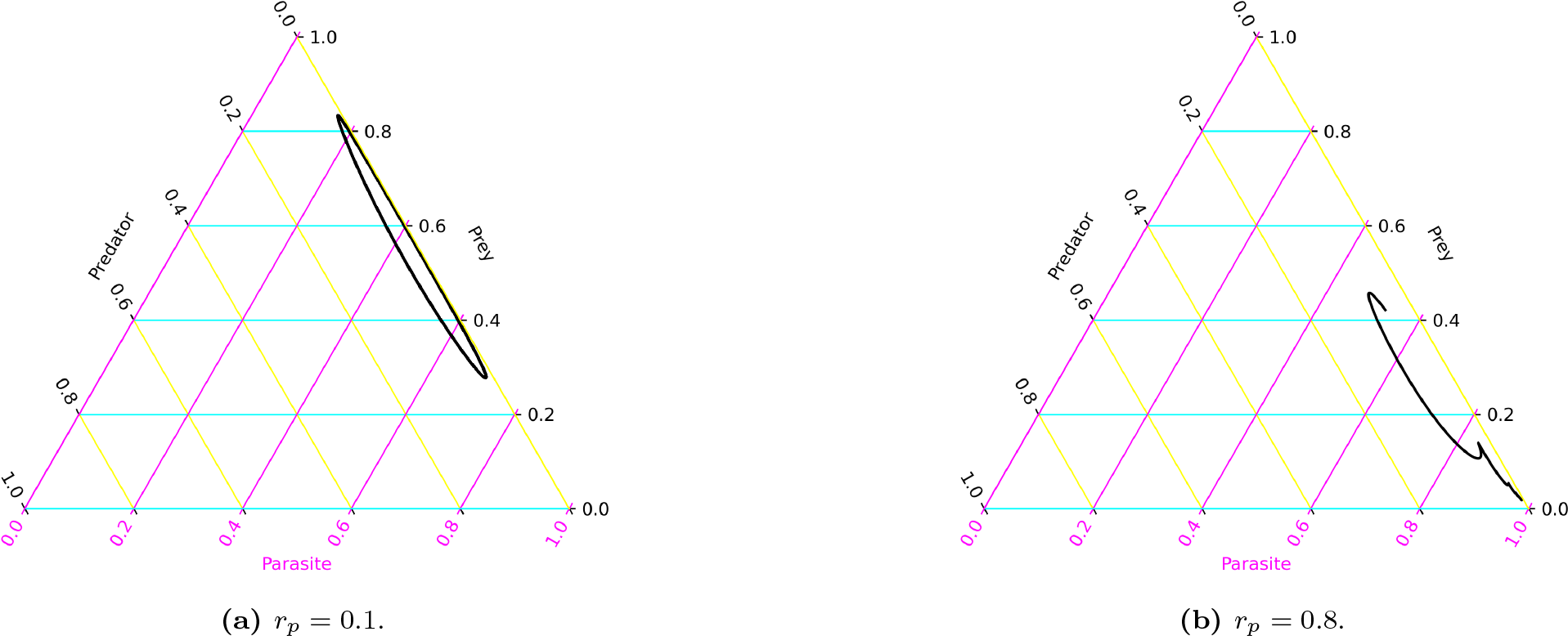
Simplex of species, according to numerical approximation of deterministic equations. Parameter values: *g*_*x*_ = 2, *d*_*x*_ = 0.1, *K* = 2000, *S* = 0.0005, *n*_*z*_ = 6, *d*_*z*_ = 0.09, *f*_*y*_ = 0.01, *k*_*y*_ = 0.2, *d*_*y*_ = 1, *r*_*x*_ = 1, *r*_*e*_ = 1, *Q*_*x*_ = 1 and *Q*_*y*_ = 0.8. Initial conditions: uninfected predator = 100, uninfected prey = 800, infected predator = 0, infected prey = 0 and parasite = 1000.

### 3 Other supplementary figures

**Figure 7:**
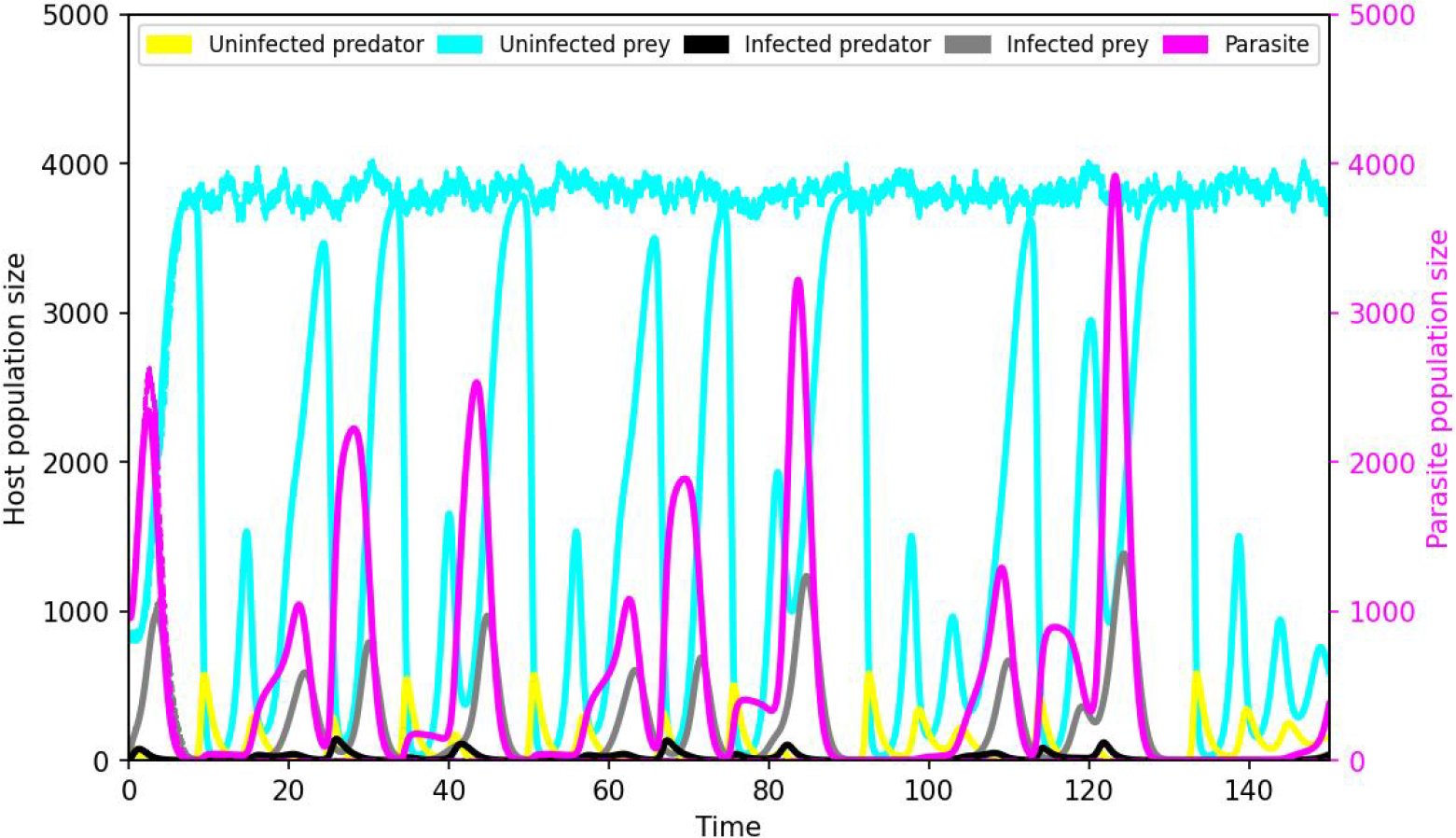
An example of the extinction of the predator species followed by the parasite species. Under a high reproduction cost of infected predators, the predator species is very likely to go extinction in stochastic dynamics. Here, we show population dynamics of uninfected predator (i.e., yellow line), uninfected prey (i.e., cyan line), infected predator (i.e., black line), infected prey (i.e., grey line), and parasites (i.e., magenta line) under both deterministic (solid lines) and stochastic dynamics (dashed lines). While the reproduction cost is so high for infected predators, the predator population (yellow and black lines) cycles to extremely low values deterministically. Correspondingly, the chance of extinction increases largely in stochastic processes due to the small population size. Thus, in this example, we do not see any dashed lines later for the predator and parasite species, and the prey species returns to a fluctuation around its own carrying capacity. Parameter values: *g*_*x*_ = 2, *r*_*x*_ = 1, *d*_*x*_ = 0.1, *K* = 2000, *S* = 0.0005, *n*_*z*_ = 6, *d*_*z*_ = 0.09, *f*_*y*_ = 0.01, *k*_*y*_ = 0.2, *d*_*y*_ = 1, *r*_*e*_ = 1, *r*_*p*_ = 0, *r*_*x*_ = 1, *Q*_*y*_ = 1 and *Q*_*x*_ = 1. Initial conditions: uninfected predator = 100, uninfected prey = 800, infected predator = 0, infected prey = 0 and parasite = 1000.

**Figure 8:**
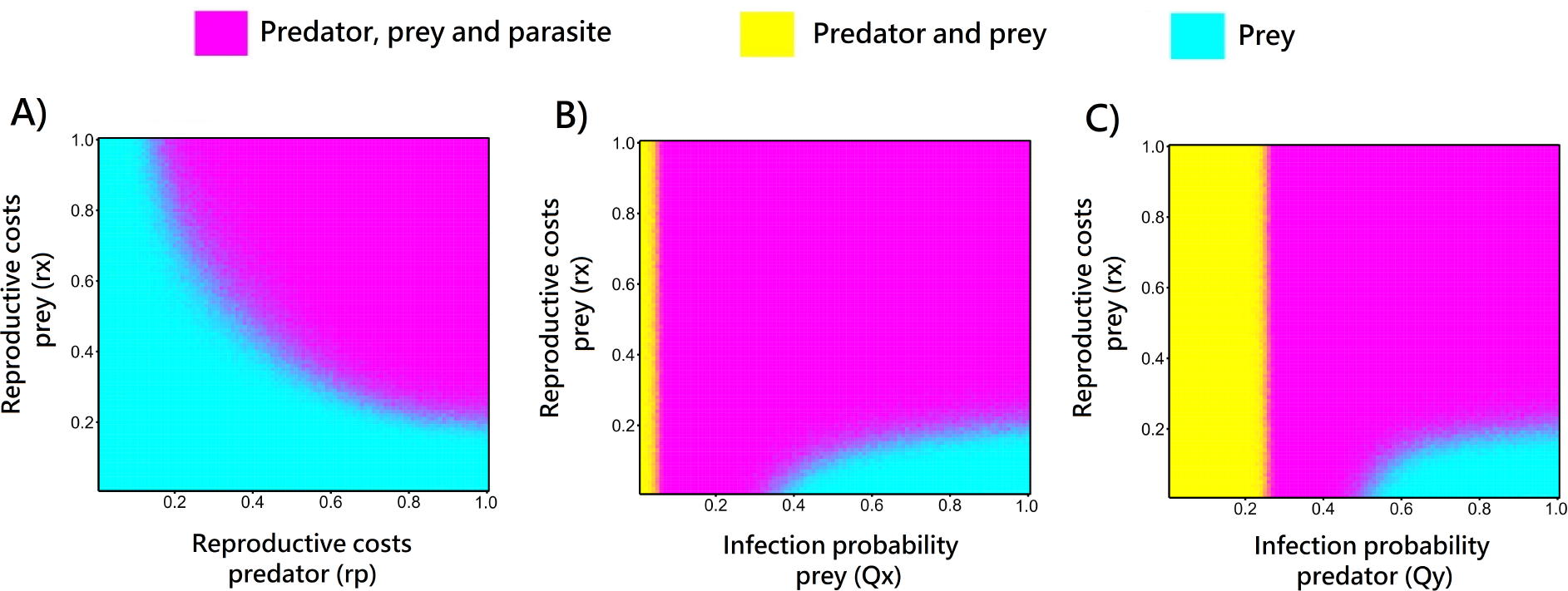
Patterns of species coexistence in stochastic simulations. Each parameter combination displays the average value from 100 independent realizations over a time period of 130-150 time steps for each run. Parameter values: *g*_*x*_ = 2, *d*_*x*_ = 0.1, *K* = 2000, *S* = 0.0005, *n*_*z*_ = 6, *d*_*z*_ = 0.09, *f*_*y*_ = 0.01, *k*_*y*_ = 0.2, *d*_*y*_ = 1, *r*_*e*_ = 1. Panel A shows *Q*_*x*_ = 1 and *Q*_*y*_ = 1; panel B shows *r*_*p*_ = 1 and *Q*_*y*_ = 1; panel C shows *r*_*p*_ = 1 and *Q*_*x*_ = 1. Initial conditions: uninfected predator = 100, uninfected prey = 800, infected predator = 0, infected prey = 0 and parasite = 1000.

## Notes

### Competing Interest Statement

The authors have declared no competing interest.

